# Genomic Characterization of *SiGRFs* in Foxtail Millet and *SiGRF1-*overexpression in *Arabidopsis thaliana* Promotes Plant to Avoid Salt Stress

**DOI:** 10.1101/849398

**Authors:** Jiaming Liu, Chengyao Jiang, Lu Kang, Chongchang Zhang, Yu Song, Weijun Zheng

## Abstract

In plants, 14-3-3 proteins are recognized as mediators of signal transduction and function in both development and stress response. However, their functions have not been reported in the C_4_ crop foxtail millet. Here, phylogenetic analysis categorized foxtail millet 14-3-3s (SiGRFs) into ten discrete groups (Clusters I to □). Transcriptome and qPCR analyses showed that all the *SiGRFs* responded to at least one abiotic stress. All but one *SiGRF-*overexpressing (OE) *Arabidopsis thaliana* line (*SiGRF1*) exhibited insensitivity to abiotic stresses during seed germination and seedling growth. Compared with the Col-0 wild-type, *SiGRF1-OEs* had slightly lower germination rates and smaller leaves. However, flowering time of *SiGRF1-OEs* occurred earlier than that of Col-0 under high-salt stress. Interaction of SiGRF1 with a foxtail millet E3 ubiquitin-protein ligase (SiRNF1/2) indicates that the proteinase system might hydrolyse SiGRF1. Further investigation showed that SiGRF1 localized in the cytoplasm, and its gene was ubiquitously expressed in various tissues throughout various developmental stages. Additionally, flowering-related genes, *WRKY71*, *FLOWERING LOCUS T*, *LEAFY* and *FRUITFULL*, in *SiGRF1-OEs* exhibited considerably higher expression levels than those in Col-0 under salinity-stressed conditions. Results suggest that *SiGRF1* hastens flowering, thereby providing a means for foxtail millet to complete its life cycle and avoid further salt stress.

**Highlight:** *SiGRFs* in foxtail millet: *SiGRF1* hastens flowering in transgenic *Arabidopsis thaliana* exposed to salt stress

## Introduction

Foxtail millet (*Setaria italica*) has been regarded as an important dietary staple in China for many millennia (Zhang et al., 2012; Liu et al., 2016a). As a C_4_ cereal crop, it not only possesses excellent drought tolerance, but also possesses an extensive germplasm collection available for research (Doust et al., 2009; Lata et al., 2013). These features accentuate this crop as a prominent genetic model for use in the study of the evolution and physiology of C_4_ photosynthesis and abiotic stress-tolerance mechanisms, particularly in response to salinity and dehydration stress (Lata et al., 2013; Liu et al., 2016a).

The 14-3-3 proteins make up a large multigenic family of regulatory proteins that are ubiquitously present in all eukaryotes (Kumar et al., 2015). They usually regulate plant development and defense from stress through protein-protein interactions by binding to target proteins containing well-defined phosphothreonine (pThr) or phosphoserine (pSer) motifs (Muslin et al., 1996; Yaffe et al., 1997; Li and Dhaubhadel, 2011). These 14-3-3 proteins interact as a dimer with a native dimeric size of ~60 kDa where each monomer in the dimer can interact with separate target proteins (Li and Dhaubhadel, 2011). This facilitates a 14-3-3 dimer to act as a scaffolding protein to induce a variety of physiological changes in the target protein (Gokirmak et al., 2010). Published research show that 14-3-3s play crucial regulatory roles in abiotic stress response pathways and ABA signaling in plants (Chen et al., 2006; Xu and Shi, 2006; Viso et al., 2007; Yohei et al., 2007; Caroline et al., 2010; Schoonheim et al., 2010; Denison et al., 2011; Vysotskii et al., 2013; Sun et al., 2015; Chen et al., 2017). In response to abiotic stress, four rice 14-3-3 genes, *GF14b*, *GF14c*, *GF14e* and *Gf14f*, were induced by the defense compounds, benzothiadiazole, methyl jasmonate, ethephon, and hydrogen peroxide. They were also differentially regulated after exposure to salinity, drought, wounding and ABA (Chen et al., 2006). The 14-3-3 proteins 14-3-3κ and 14-3-3χ can undergo self-phosphorylation by stress-activated kinases, such as SnRK2.8 in *A. thaliana*. In relation to ABA signaling, 14-3-3 proteins have been shown to be present at the promoters of two *A. thaliana* late-embryogenesis genes, *AtEm1* and *AtEm6*, which are induced by *ABI3*, an ABA-regulated transcription factor (Viso et al., 2007). Five 14-3-3 isoforms interact with the ABA-regulated transcription factors and are involved in ABA signal transduction during barley seed germination (Schoonheim et al., 2010). A soybean 14-3-3 protein can regulate transgenic *A. thaliana* ABA sensitivity (Sun et al., 2015). Rice OsCPK21 phosphorylates 14-3-3 proteins in response to ABA signaling and salt stress (Chen et al., 2017).

Salt stress is a major abiotic stress in the production of foxtail millet. The 14-3-3 proteins described above are involved in salt-stress response in C3 plants, but there is no comprehensive genome-wide research of 14-3-3 proteins and abiotic stress in foxtail millet. In this study, a comprehensive *in silico* expression analysis of the 14-3-3 genes, hereafter called *SiGRF* (GENERAL REGULATORY FACTOR) genes for simplicity, at several development stages of foxtail millet were performed using available microarray data. The expression patterns under stress conditions showed that all the *SiGRFs* were responsive to at least one abiotic stress. Phenotypic identification of overexpression of *SiGRFs* in *A. thaliana* further confirmed the stress resistant functions of the *SiGRF*s. In addition, we studied the function of *SiGRF1* in detail, including gene expression pattern, protein subcellular localization, candidate interaction protein screening, protein-protein interaction verification, and phenotypic characteristics of *SiGRF1*-*OE*s under salt stress. Results imply that SiGRF1 hastens transgenic *A. thaliana* flowering in the presence of salt stress to achieve reproduction despite the harsh environment.

## Materials and methods

### Plant materials and stress treatments of foxtail millet

Foxtail millet Yugu 1, known for its tolerance to abiotic stress, was used to amplify cDNA sequences of *SiGRFs*, the *SiGRF1* gene promoter, and *SiRNF1/2* (Si021868m). *Arabidopsis thaliana* Columbia-0 (WT) was used as the background for overexpressing *SiGRFs.* Foxtail millet seeds were germinated and grown in a growth chamber at 28 ± 1°C day and 23 ± 1°C night temperatures with 70 ± 5 % relative humidity and a photoperiod of 14 h. *Arabidopsis thaliana* seeds were synchronized on wet filter paper at 4 °C for 3 d and then sown in soil. They were kept in a greenhouse at 20–22 °C with 45% relative humidity under long-day (LD) conditions (16 h light/8 h dark). The stress treatments were performed as previously described (Liu et al., 2016a).

### Identification of 14-3-3 genes and evolutionary analyses

The hidden Markov model (HMM) profile of the 14-3-3 domain (PF00244) downloaded from Pfam v27.0 (http://Pfam.sanger.ac.uk/) was used to identify 14-3-3 proteins in *A. thaliana* (GRF), *Brachypodium distachyon*, *Oryza sativa*, *Triticum aestivum*, *Sorghum bicolor*, maize, and foxtail millet as described in a previous report (Liu et al., 2016a). The amino acid sequences of the seven species’ 14-3-3 proteins were obtained using BLASTP in the protein database of the National Center for Biotechnology Information (http://blast.ncbi.nlm.nih.gov/Blast.cgi). Sequence alignment was performed by ClustalX (Thompson et al., 1997). The complete amino acid sequences of 14-3-3 proteins were used and the neighbor-joining method was adopted to construct a phylogenetic tree by the MEGA5.1 program, and the confidence levels of monophyletic groups were estimated using a bootstrap analysis of 500 replicates (Tamura et al., 2011). The full-length open reading frames of *SiGRFs* and *SiRNF1/2* were obtained from foxtail millet cDNA. The primers for cloning are listed in Supplemental Table 1. The PCR products were cloned into pLB vectors (TianGen, China) and sequenced with an ABI 3730XL 96-capillary DNA analyzer (Lifetech, USA).

### Transcriptome analysis

The gene transcriptome data for foxtail millet were obtained from Phytozome v.12.1 (https://phytozome.jgi.doe.gov/pz/portal.html). Data from the following foxtail millet tissue/organs and developmental stages (for some tissues) were analyzed: etiolated seedling, root, shoot, leaves (1 to 6 from bottom to top of the shoot), and panicles (collected from the 5th and 10th day after heading) (https://phytozome.jgi.doe.gov/pz/portal.html). Tissues samples were collected from plants exposed to various light- or nitrogen-source treatments according to a foxtail millet study (B100) described in Phytozome v.12.1. For the light experiments, plants were grown under continuous monochromatic light (blue: 6 µmol m^−2^ s^−1^, red: 50 µmol m^−2^ s^−1^, or far-red: 80 µmol m^−2^ s^−1^), and watered with RO water every three days. Total aerial tissues were collected (at 9:30 am) from 8-day-old seedlings. For the nitrogen treatments, plants were grown for 30 days under differing nitrogen source regimes, 10 mM KNO_3_ (NO_3_-plants), 10 mM (NH_4_)_3_PO_4_ (NH_4+_ plants) or 10 mM urea (urea plants). Root tissue was harvested from the nitrogen-treated plants.

### DNA isolation, RNA extraction, RT-PCR and qRT-PCR

Genomic DNA for each sample was isolated from foxtail millet leaves using the CTAB method (Saghai-Maroof et al., 1984). Isolation of total RNA from plant materials was performed using an RNA extraction kit (Takara, Japan) according to the manufacturer’s recommendations. Synthesis of cDNA and RT-PCR were conducted as previously described (Liu et al., 2016a). Analysis by qPCR was carried out with TransScript® II Probe One-Step qPCR SuperMix (TransGen, China) on an ABI 7500 system. The foxtail millet *SiActin* gene (Si036655m) was used as an internal control to normalize all data. The 2 ^−△△CT^ method (Livak and Schmittgen, 2001) was used to evaluate the relative expression of each gene. All RT-PCR reactions were repeated three times.

### Subcellular localization

The coding region of *SiGRF1* without the termination codon was inserted at the BamH I site of the subcellular localization vector p16318, which contained the 35S promoter and C-terminal green fluorescent protein (GFP) (Liu et al., 2016a; Liu et al., 2016b). The transient expression assays were performed as previously described (Liu et al., 2016a), and then samples were observed at 488 and 543 nm illumination using a Zeiss LSM700 microscope.

### GUS histochemical assay

A 2.5-kb *SiGRF1* promoter region was inserted into the vector pCAMBIA1305 at the EcoR I site. The construct was introduced into *A. thaliana* Columbia-0 by *Agrobacterium tumefaciens*-mediated transformation as described previously (Jeon et al., 2000). Tissues from individuals of the homozygous T_3_ generation of transgenic *pSiGRF1::GUS A. thaliana* were sampled at different growth stages and stained in a GUS staining solution as previously described (Ma et al., 2015). Samples were destained in 50, 70, and 100 % ethanol for 5 min, consecutively, and then bleached by immersion in 100 % ethanol. The decolorized tissues were observed by bright field microscopy (LEICA M165FC, Germany) and photographed using a digital camera (LEICA DFC420C, Germany).

### Generation of transgenic *A. thaliana*

The coding sequences of *SiGRF* genes were amplified and cloned into pCAMBIA1305-GFP under the control of the *cauliflower mosaic virus* (CaMV) 35S promoter, resulting in a *35S::SiGRFs* construct. The constructs were confirmed by sequencing and then transformed into WT *A. thaliana* of the ecotype Col-0. Lines of the *35S::SiGRFs* transgenics, with various expression levels of the *SiGRFs* gene, were obtained for further analysis.

### Seed germination and root growth assays

For the germination assay, seeds were subjected to 100 mM NaCl, 6% (w/v) polyethylene glycol (PEG) 6000 (to simulate osmotic stress), or 0.5 μM ABA treatments. For the root growth assays, 5-day-old seedlings were grown on vertical agar plates in the presence or absence of 100 mM NaCl, 6% (w/v) PEG 6000, and 5 μM ABA. Root lengths were measured after five days of treatment. The root lengths and lateral root numbers of plants were monitored and data were statistically analyzed. For the germination assay of *SiGRF* genes in transgenic plants, the seed number was recorded every 12 h post-incubation for visible radical emergence as a proxy for seed germination. Each treatment contained three independent replicates.

### Yeast two-hybrid assay

SiGRF1 protein interactions were investigated by screening a foxtail millet cDNA library in yeast with the Matchmaker Two-Hybrid System 3 (Clontech). *SiGRF1* cDNA without the termination codon was cloned with EcoR1 technology in pGBKT7. The bait vector obtained was transformed in the yeast strain AH109. The bait strain was transformed according to the manufacturer’s instructions with a commercially available library (Takara, Japan) expressing Nub fused to cDNAs. The library was constructed with total RNA from a mixture of tissues from 10-d-old foxtail millet seedlings.

### Bimolecular fluorescence complementation (BiFC)

The coding sequences of *SiGRF1* was inserted into the BamH I sites of the pSPYNE vector and that of *SiRNF1/2* was inserted into pSPYCE vectors (Walter et al., 2004). These plasmids were extracted by QIAGEN Plasmid Maxi Kit and transformed into wheat leaf protoplasts. The YFP fluorescence of protoplasts was assayed under a Zeiss LSM 700 confocal microscope 8 h after transformation.

### Co-immunoprecipitation (Co-IP) assays

To evaluate the interaction between SiGRF1 and SiRNF1/2 *in vivo*, Co-IP assays were carried out as previously described (Zheng et al., 2013). *Agrobacterium* strain GV3101 carrying the pCAMBIA1305-SiGRF1-GFP, pCAMBIA1305-SiRNF1/2-FLAG, or 35S::p19 construct was co-infiltrated into a *Nicotiana benthamiana* leaf. After growing in the dark for three days, the leaf was harvested and native extraction buffer (50 mM Tris-MES [pH 8.0], 0.5 M sucrose, 1 mM MgCl_2_, 10 mM EDTA, 5 mM DTT, and 1× protease inhibitor cocktail) was used to lyse the sample. Then, 250 μL of extract was incubated with 20 μL anti-GFP conjugated agarose (Sigma-Aldrich) for 8 h at 4°C. The agarose was washed twice with 200 μL native extraction buffer. The pellet was detected by immunoblot analysis for anti-FLAG and anti-GFP.

## Results

### Phylogenetic Analysis

The foxtail millet genome contains eight *SiGRF*s, and the predicted polypeptide lengths of SiGRF proteins ranged from 429 to 731. These protein sequences have small variations in both isoelectric point (p*I*) values (ranging from 4.71 to 5.19) and molecular weights (ranging from 26.188 to 29.690 kDa; Table S2).

To evaluate the phylogenetic relationships among the *SiGRF*s in foxtail millet and six other species, predicted 14-3-3 sequences of 15 *A. thaliana*, 16 *B. distachyon*, 10 *O. sativa*, 28 *T. aestivum*, 6 *S. bicolor*, 26 maize, and 8 foxtail millet were used to generate a neighbor-joining phylogenetic tree (Figure 1). The phylogenetic analysis categorized the 14-3-3s into ten discrete groups (Clusters I to □), containing, respectively, 20, 14, 7, 4, 17, 3, 8, 14, 10, and 11 predicted proteins (Figure 1). Many of the internal branches had high bootstrap values, indicating statistically reliable pairs of possible homologous proteins.

**Figure 1.**
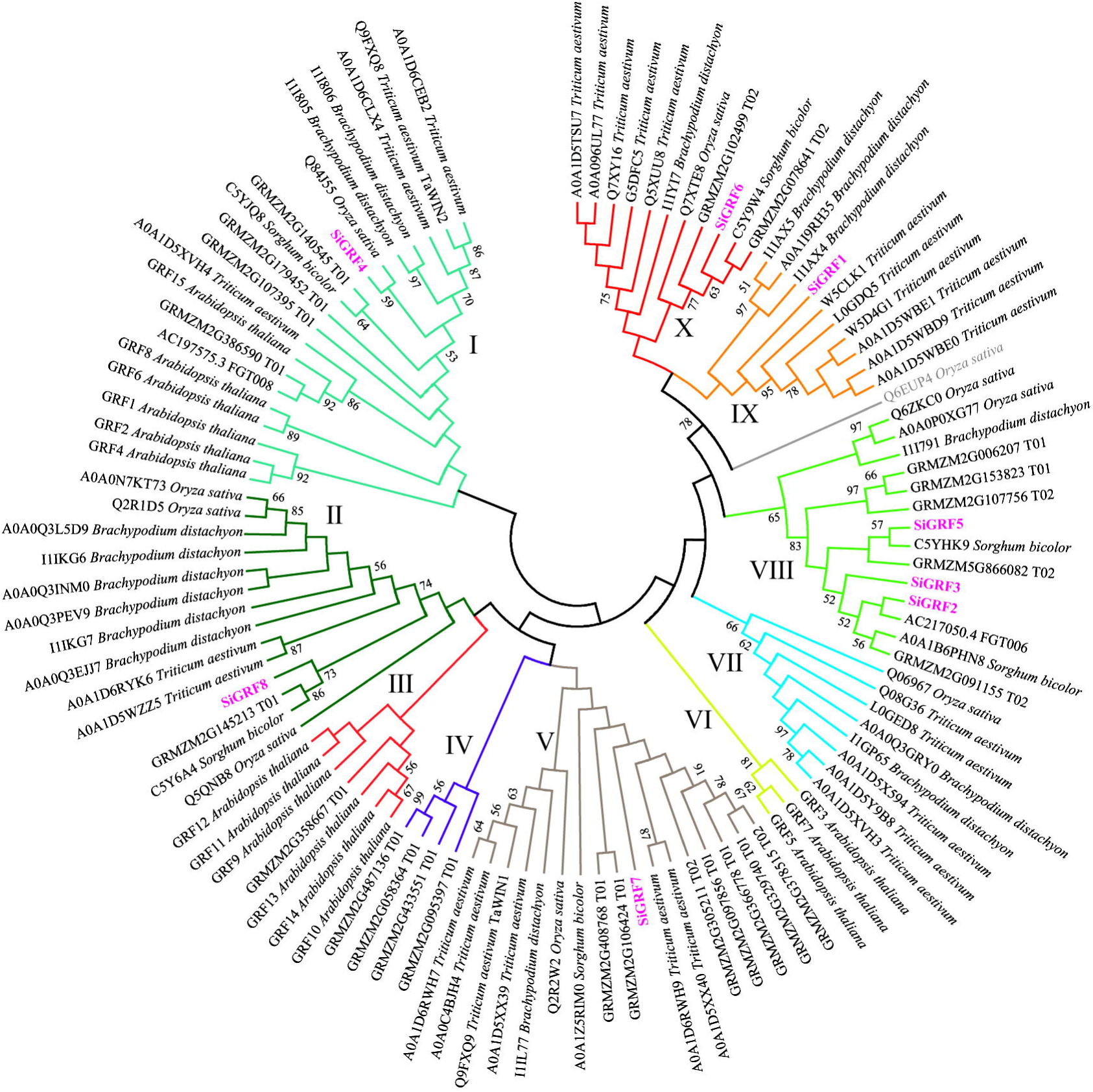
Neighbor-joining distance tree of 14-3-3 proteins in seven species. The tree is presented radially so that distances from the center represent cumulative branch lengths. Terminal branches and labels are colored to indicate different groups and foxtail millet (pink).

### *In silico* tissue-specific expression profiling of *SiGRF*s

A heat map generated for examining tissue-specific expression showed differential transcript abundances of eight *SiGRF* genes in four major tissues, namely shoot (control), germ shoot, leaf, and panicle (Figure 2). The average log signal values for all of the *SiGRF* genes from three biological replicates of each sample are given in Table S3. The results showed greater levels of expression in all the plant tissues compared to that of the shoot. Greater expression of *SiGRF1* and *SiGRF2* were only observed in the germ shoot. Lower levels of *SiGRF5*, *SiGRF7* and *SiGRF8* resulted in the leaf (Figure 2). *SiGRF8* was stronger in both panicle stages than in the shoot (Figure 2).

**Figure 2.**
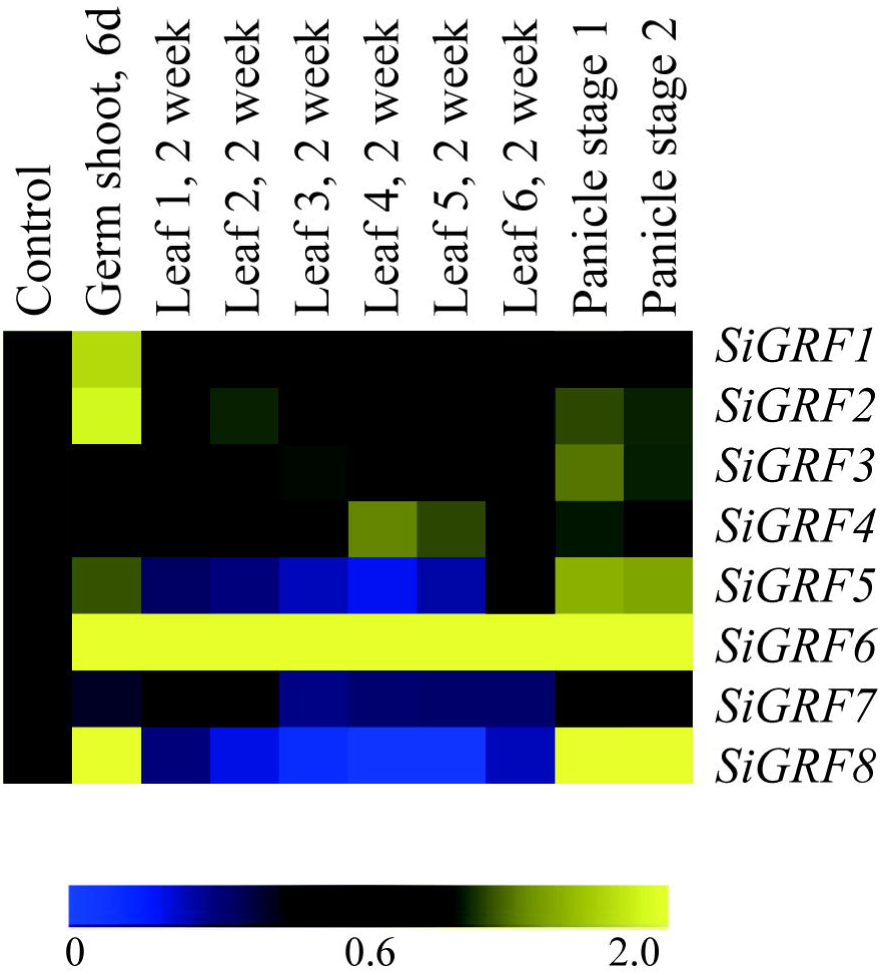
Tissue-specific expression of foxtail millet *SiGRF* genes in four major tissues: shoot (control), germ shoot, leaf, and panicle (two stages). The color scale representing the log of signal values is shown below the expression profiles.

### Expression analysis of *SiGRF*s under abiotic stress and light conditions

To explore the potential functions of *SiGRF* genes under different stress and light treatments in foxtail millet, microarray analysis was performed using available GeneATLAS data (https://phytozome.jgi.doe.gov/pz/portal.html) for RNA from foxtail millet roots subjected to ammonia, drought, nitrate, or urea and shoots exposed to dark, blue, red, or far red light treatments. Results showed that all of the *SiGRF* genes varied in their expression levels in response to one or more stress or light relative to their expression in untreated control samples (Figure 3 and Table S4). Expression levels of *SiGRF4* and *SiGRF8* were stronger than that of their respective controls across most treatments. Compared with the respective control, *SiGRF2*, *SiGRF3*, *SiGRF5*, and *SiGRF7* in aerial tissues were lower in expression under blue light and far red light treatments (Figure 3), which suggests that these genes may be related to plant growth and flowering.

**Figure 3.**
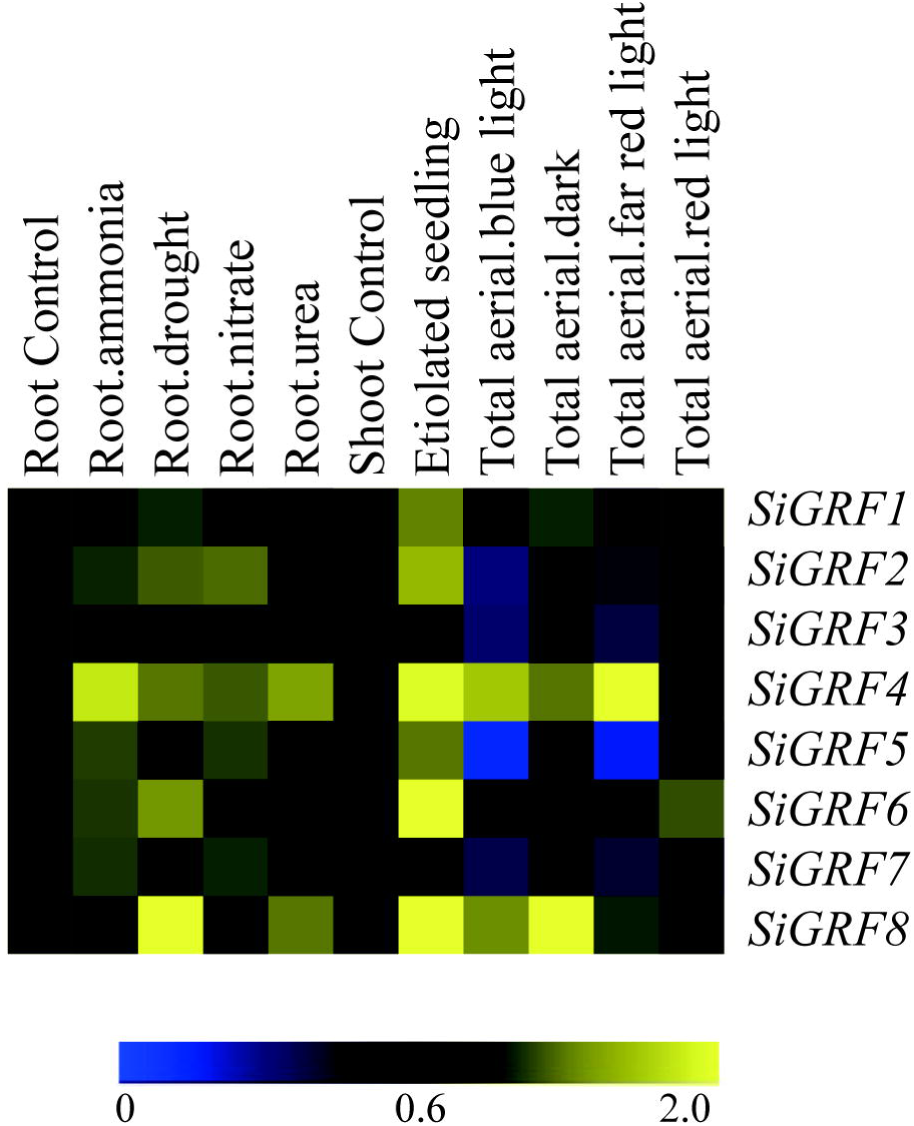
Expression profiles of *SiGRF* genes differentially expressed under stress and light conditions. The values of *SiGRF* genes under the control (untreated) and various stress conditions (labeled at the top of each lane) are presented by cluster display. The color scale representing signal values is shown at the bottom.

### Verification of microarray data using qPCR

We used qPCR to further verify the expression levels of *SiGRF* genes under abiotic stress or exogenous ABA (Figure 4). The results show that the expression levels of four genes were up-regulated (> 2 fold) by salt stress, four were up-regulated by 6% PEG, and five were up-regulated by ABA. The expression of *SiGRF1* was up-regulated under both salt and ABA stress treatments. *SiGRF6* and *SiGRF8* expression levels were up-regulated under ABA and 6% PEG stress conditions, respectively. Notably, the expression of *SiGRF2*, *SiGRF4* and *SiGRF7* were up-regulated under salt, 6% PEG, and ABA stress conditions. The expression levels of two *SiGRF* genes (*SiGRF3* and *SiGRF5*) were unchanged by any treatment.

**Figure 4.**
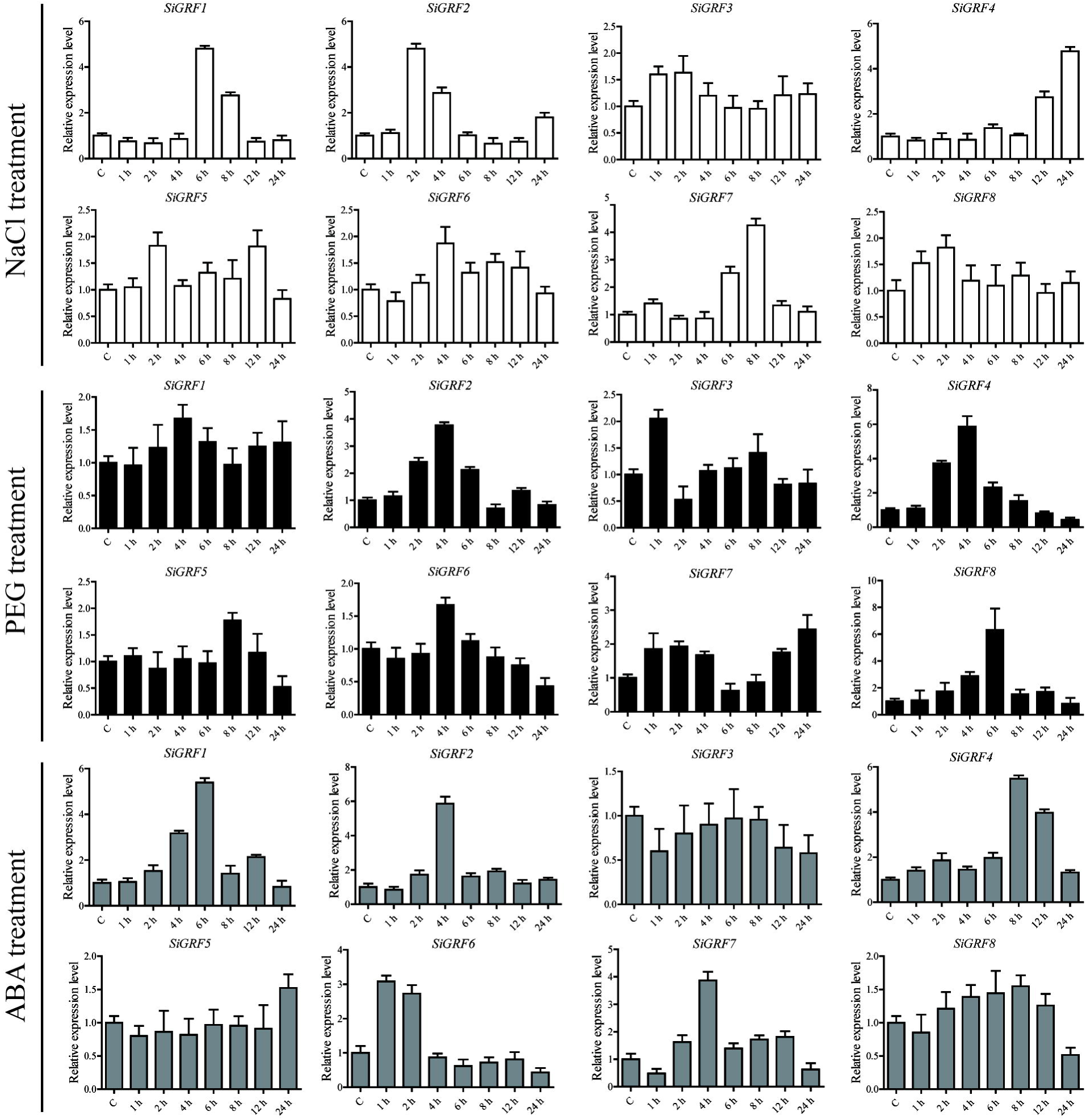
Relative expression levels of *SiGRF* genes analyzed using qPCR under treatment of salinity stress, 6% PEG, or ABA for 1, 2, 4, 6, 8, 12, or 24 h. The relative expression level of each gene was calculated relative to the expression in the respective untreated control samples (0 h). Foxtail millet *SiActin* (Si036655m) was used as an internal control to normalize the expression data. The error bars represent the standard deviation calculated based on three technical replicates for each of the biological duplicates.

### Phenotypic analysis of *SiGRF*-transgenic *A. thaliana*

The full-length open reading frames of eight *SiGRF*s were obtained from foxtail millet cDNA and introduced into *A. thaliana* Col-0 to generate several *35S::SiGRF* lines for each gene. Four overexpression lines of each *SiGRF* gene were selected for stress tolerance assays. Due to space limitations, we only show two lines (Figure S1–4). Under the control conditions, we observed no significant differences in the growth or morphology between *SiGRF-*OEs and Col-0 plants, with the exception of *SiGRF1-*OEs. The seed germination of *SiGRF2-*OEs to *SiGRF7-*OEs displayed a phenotype indistinguishable from that of Col-0 under the control condition (Figure S1 and S2). However, the germination of *SiGRF5-*OE, *SiGRF6-*OE, and *SiGRF8-*OE lines was faster than that of Col-0 under 0.5 μM ABA (Figure S2). In the presence of 120 mM NaCl, *SiGRF4-*OE and *SiGRF5-*OE lines produced longer main roots, and *SiGRF7-*OE lines produced longer lateral roots compared to those of Col-0 (Figure S3 and S4). When grown on 6% PEG medium, *SiGRF2-*OEs and *SiGRF8-*OEs had longer lateral roots than that of Col-0 (Figure S3 and S4). *SiGRF2-*OE, *SiGRF4-*OE, *SiGRF6-*OE, *SiGRF7-*OE, and *SiGRF8-*OE lines exhibited longer main root lengths, and *SiGRF7-*OEs also exhibited longer lateral lengths compared to the respective root lengths of Col-0 under 0.5 μM ABA (Figure S3 and S4).

*SiGRF1-OE*s had slightly lower germination rates and smaller leaves compared to Col-0 under the control condition (Figure S1, S3, 5A and 5B). In the presence of 120 mM NaCl, 6% PEG or 0.5 μM ABA, the germination of *SiGRF1-OE*s was slower than that of Col-0, and their seedlings’ root lengths were shorter compared to that of Col-0. The lateral roots of *SiGRF1-OE*s were longer than lateral roots of Col-0 under the 120 mM NaCl treatment (Figure S1), suggesting that *SiGRF1* may function in resistance to salt stress.

### Flowering of *SiGRF1-*OEs was relatively insensitive to salt stress

Assays were performed to further characterize the function of *SiGRF1*. Under standard growth conditions, we observed no significant differences in the flowering initiation means dates and total leaf means numbers between *SiGRF1-OE*s and Col-0 plants (Figure 5 C, D and E). The flowering time of *SiGRF1-OE*s occurred earlier than that of Col-0 under high salt stress and earlier than that of untreated *SiGRF1-OE*s (Figure 5 C, D and E). We also observed that GFP fluorescence signals of *SiGRF1-OE*s increased significantly under the salt treatment compared to under the control condition (Figure 5 F). Together with the longer lateral root phenotype of *SiGRF1-OE*s under the 120 mM NaCl treatment, these results suggest that overexpression of *SiGRF1* may help transgenic *A. thaliana* complete its life cycle quickly to avoid salt stress.

**Figure 5.**
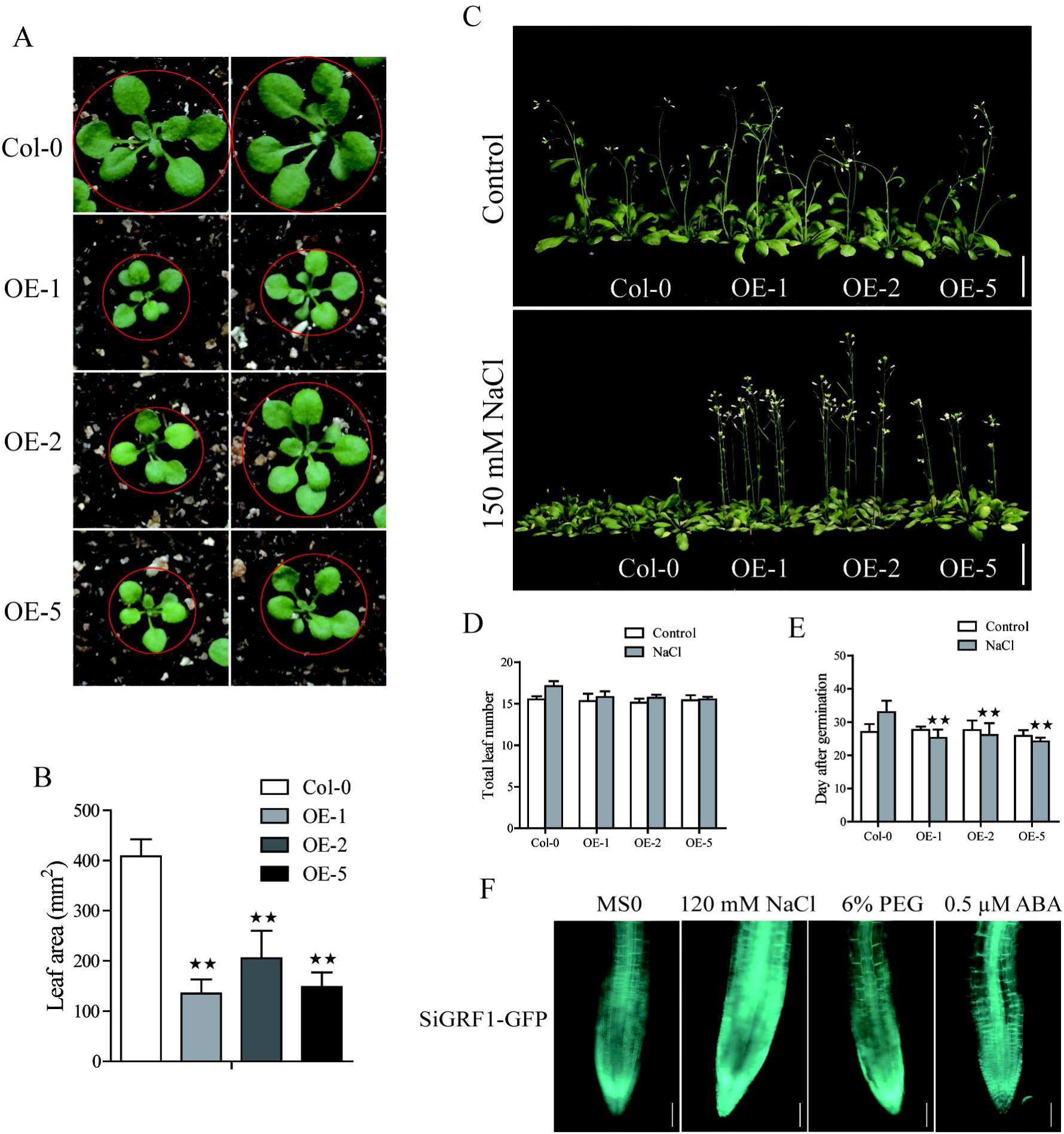
Phenotype analysis of *SiGRF1*-OEs. (A) and (B) *SiGRF1-OE*s have a smaller leaf size compared to that of the wild-type (Col-0) under the non-stressed condition. Bars = 1 cm. (C) The flowering time of *SiGRF1-OE*s under no stress and high salt stress. Bars = 5 cm. (D) and (E) Flowering days and number of leaves at flowering of *SiGRF1-OE*s under no stress and high salt stress. Data represent means ± SD (n = 100). (F) The GFP fluorescence signal of *SiGRF1-OE*s when plants were grown on MS medium with or without treatment of 120 mM NaCl, 6% PEG, or 0.5 μM ABA for 1 h. Bars = 0.1 cm.

### RNA-Seq analysis of gene expression associated with SiGRF1 action

To identify changes in gene expression associated with SiGRF1 action, we performed an RNA-seq analysis of total RNA isolated from 2-week-old seedlings, comparing *SiGRF1-OE* line 1 plants to Col-0 plants. We identified 174 genes that were differentially expressed (log2 value ≥ 1.5-fold difference, p-value less than 0.05), which comprised of 156 up-regulated genes and 18 down-regulated genes. Di□erentially expressed genes were categorized into functional groups using Gene Ontology (GO) analyses. The top key GO terms were cellular process, response to stimulus, metabolic process, and development process (Figure 6 A). Notably, the genes related to seed germination and flower development, *OLEOSIN1* (*OLEO1*), *OLEOSIN2* (*OLEO2*), *OLEOSIN4* (*OLEO4*), *AT3G01570*, *EARLY LIGHT-INDUCABLE PROTEIN* (*ELIP1*), *WRKY71*, *CRUCIFERIN 3* (*CRU3*), *AGAMOUS-like 67* (*AGL67*), *MADS AFFECTING FLOWERING 4* (*MAF4*), and *LIPID-TRANSFER PROTEIN* (*AT4G22460*), suggesting that the phenotypes of *SiGRF1-OE* lines under normal and salt stress condition may be caused by abnormal expression of these genes (Figure S3, 5 and 6B). We used qPCR to further verify the accuracy of the RNA-seq analysis. The results show that the expression levels of these genes were roughly consistent with the RNA-seq analysis (Figure 6 C). It is worth noting that the expression level of *WRKY71* was significantly greater in *SiGRF1-*OE line 1 plants than the expression level of Col-0 plants under salt stress (Figure 6 B and C), and Yu et al. (2008) has shown that *WRKY71* acts antagonistically to salt-delayed flowering in *A. thaliana*. Furthermore, because flower initiation days of *SiGRF1-OE*s noticeably occurred earlier than that of Col-0 under high salt stress (Figure 5 C), we suspect that the *SiGRF1* gene may regulate expression of *WRKY71* to induce the earlier flowering time in transgenic plants under salt stress.

**Figure 6.**
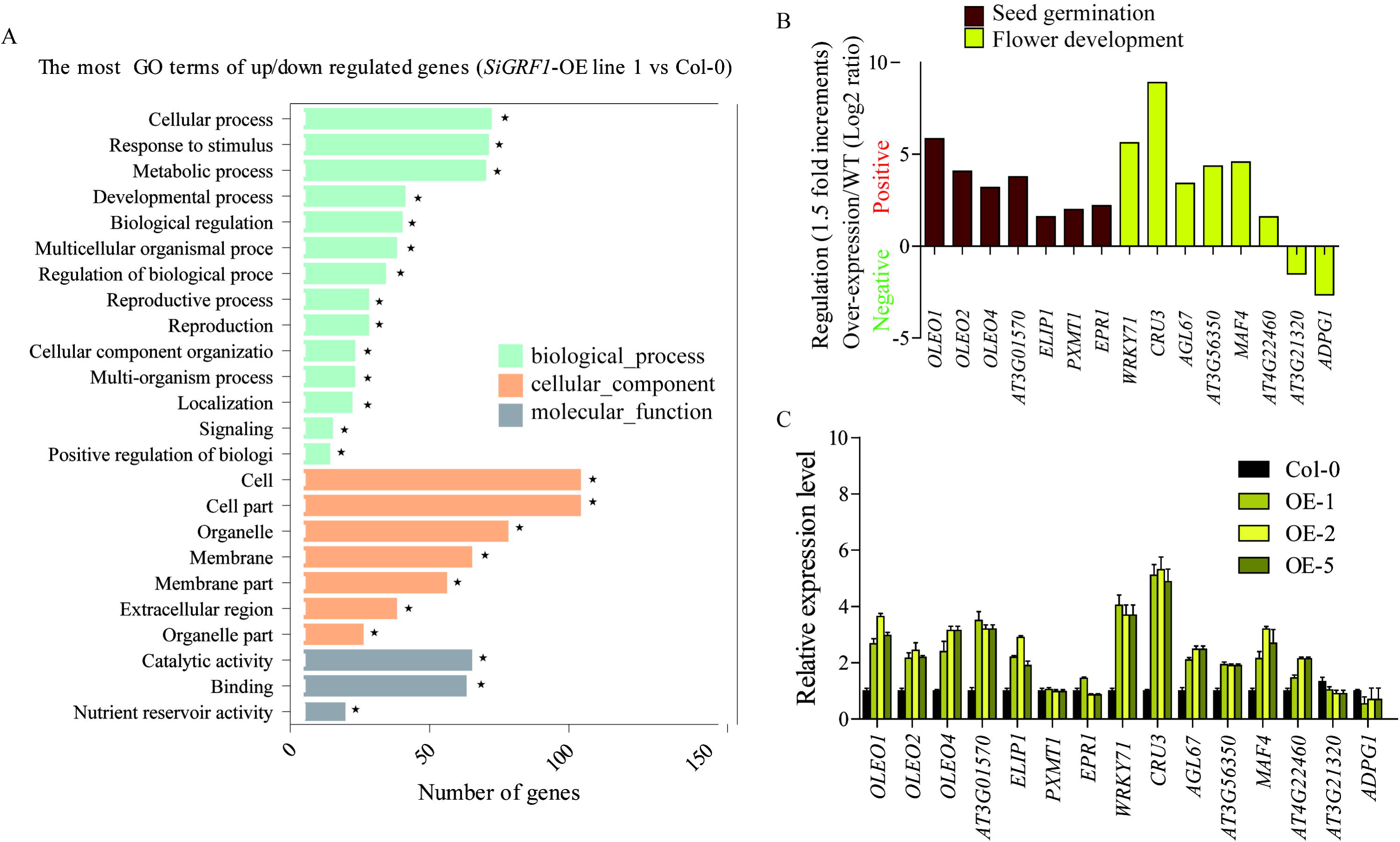
SiGRF1 activity can change the expression of genes related to seed germination, stress response and flower development. (A) The enrichment of GO terms for genes that were differentially expressed between *SiGRF1-OE* line 1 and Col-0 was analyzed using RNA-seq. Vertical coordinates represent enriched GO terms, and horizontal coordinates represent the numbers of differentially expressed genes for these GO terms. The light green columns represent the GO terms for biological processes, orange columns represent the GO terms for cellular components, and gray columns represent the GO terms for molecular functions. Asterisks indicate significantly enriched GO terms. (B) Expression levels of genes related to seed germination and flower development from RNA-seq data. The brown columns represent seed germination, and the yellow columns represent flower development. (C) qPCR verified the accuracy of the RNA-seq data. Vertical coordinates represent fold changes, and horizontal coordinates represent different genes. The relative expression levels of those genes in Col-0 were set to “1”. *Arabidopsis actin2* (AT3g18780) was used as a reference. Error bars indicate the SDs. The data represent the means ± SD of three biological replications.

We also observed that the expression of several flower-related marker genes in *SiGRF1-OE*s (Yu et al., 2018), including *FLOWERING LOCUS T* (*FT*), LEAFY (*LFY*) and *FRUITFULL* (*FUL*), were indistinguishable from those of Col-0 under the control condition (Figure 7). However, transcript levels of these genes in *SiGRF1-OE*s plants were significantly greater due to salt stress (Figure 7). Previous studies have shown that *FT* and *LFY* were the direct targets of *WRKY71*; *FUL* was also up-regulated by *WRKY71* (Yu et al., 2017). Altogether, these data also suggest that the *SiGRF1* gene may hasten transgenic *A. thaliana* to complete its life cycle sooner to avoid salt stress via regulation of the expression of *WRKY71*.

**Figure 7.**
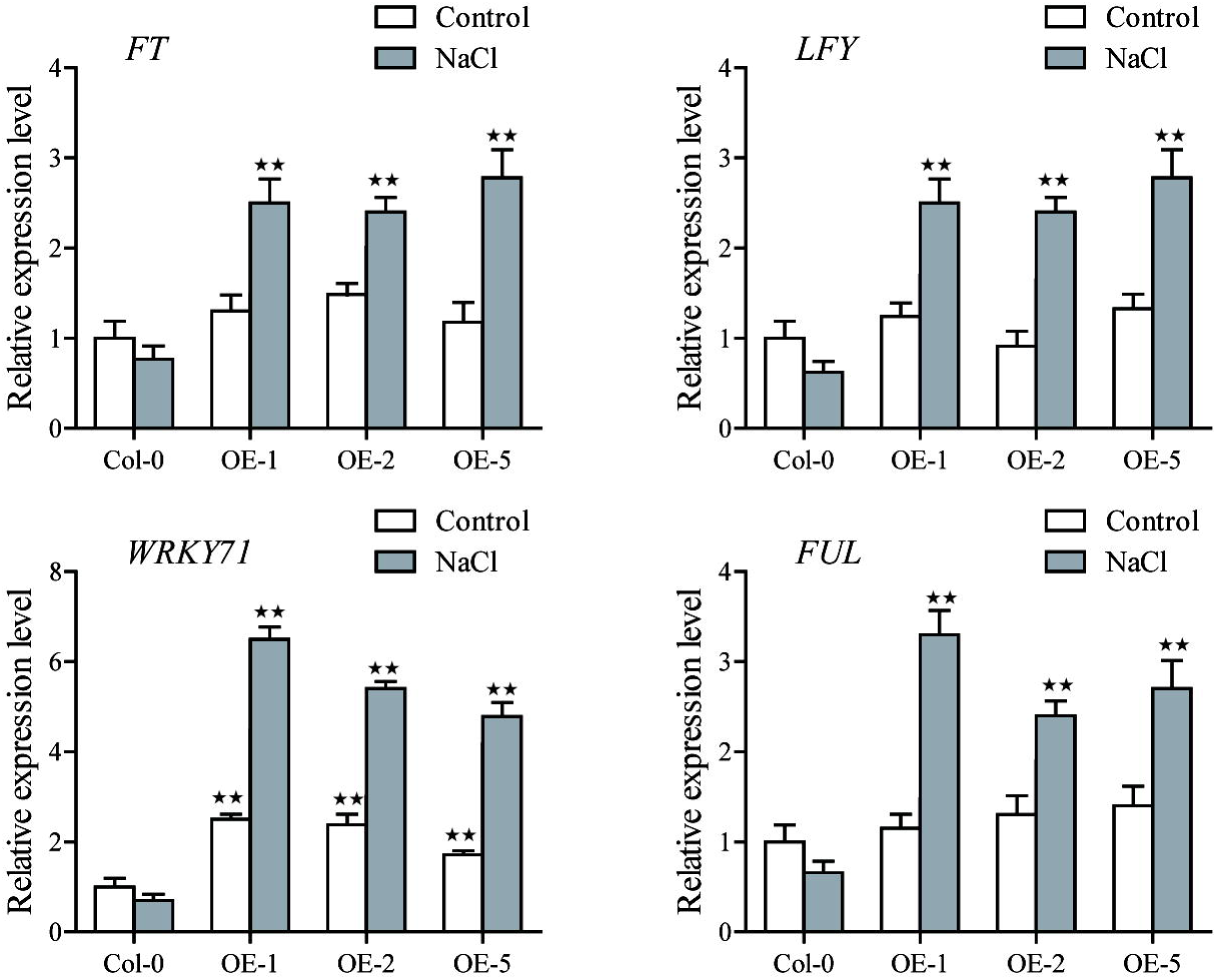
The expression levels of *FLOWERING LOCUS T* (*FT*), LEAFY (*LFY*) and *FRUITFULL* (*FUL*) in Col-0 and *SiGRF1-OE*s under no stress or salt stress conditions. Values obtained by qPCR represent means ± SD from three biological replicates.

### Identification of SiGRF1 target proteins using the yeast two-hybrid system

The identification of protein partners should provide clues to understanding the function(s) of SiGRF1. To this end, we sought to identify possible SiGRF1 target proteins using a yeast two-hybrid approach. SiGRF1, as bait protein, was used to screen the foxtail millet cDNA library. Of ~9×10^7^ primary transformants, 200 HIS-selected clones that showed LacZ activity were obtained. From these, 60 clones were randomly chosen and further analyzed by DNA sequencing. Twelve different cDNA clones contained sequences of *SiRNF1/2* (Si021868m), a member of the ubiquitin–proteasome system encoding a E3 ubiquitin-protein ligase. (Sadanandom et al., 2012) reported that cells can respond quickly to intracellular signals and varying environmental conditions because the ubiquitin–proteasome system was involved in many cell physiological processes in the removal of abnormal peptides and short-lived cytokines. The results prompted us to speculate whether or not SiGRF1 interacts physically with SiRNF1/2 and SiGRF1 might be hydrolyzed by the proteinase system.

### SiGRF1 interacted with SiRNF1/2

We next investigated the protein interactions between SiGRF1 and SiRNF1/2; the protein interactions observed in yeast were further investigated *in planta* (Figure 8 A). For this, we used the BiFC analysis to confirm whether SiGRF1 associates with SiRNF1/2 *in vivo*. Constructs for expression of the fusion proteins SiGRF1-YNE and SiRNF1/2-YCE were transiently expressed in the protoplasts from foxtail millet seedlings. Different combinations of TaPI4KIIγ-YNE/TaUFD1-YCE were used as positive controls because they have been shown to interact with each other in the plasma membrane (Liu et al., 2013), and GFP alone was the blank control. SiGRF1-YNE/SiRNF1/2-YCE and YNE/SiRNF1/2-YCE were co-transformed and the protoplasts were observed under a confocal microscope to detect yellow fluorescent protein (YFP) signals. The results showed that GFP alone was expressed throughout the cell, and TaPI4KIIγ-YNE interacted with TaUFD1-YCE in the plasma membrane (Figure 8 B and C), which are consistent with results of previous research (Liu et al., 2013). A strong YFP signal was observed in the cytoplasm of the protoplast co-transformed with SiGRF1-YNE/SiRNF1/2-YCE plasmids (Figure 8 D), whereas no YFP signal was detected in the absence of SiGRF1 (Figure 8 E). These results suggest that the interactions between SiGRF1 and SiRNF1/2 occurred in the cytoplasm (Figure 8 D). Subsequently, we performed co-immunoprecipitation assays in *N. benthamiana* leaf to further investigate whether SiGRF1 interacts with SiRNF1/2 *in vivo*. Green fluorescent protein and FLAG tags were translationally fused to the C-terminus of SiGRF1 and SiRNF1/2, respectively. *Agrobacterium* strain GV3101 carrying the *35S::SiGRF1*-GFP, *35S::SiRNF1/2*-FLAG, or *35S::p19* construct was co-infiltrated into a *N. benthamiana* leaf. Total protein was used for immunoblot analysis with anti-GFP and anti-FLAG antibodies (Figure 8 F). Anti-GFP affinity gels were used to perform immunoprecipitation. After washing, immunoblots were probed with an anti-FLAG antibody. SiRNF1/2-FLAG was pulled-down using SiGRF1-GFP. These results showed that SiGRF1 interacts physically with SiRNF1/2 in the cytoplasm.

**Figure 8.**
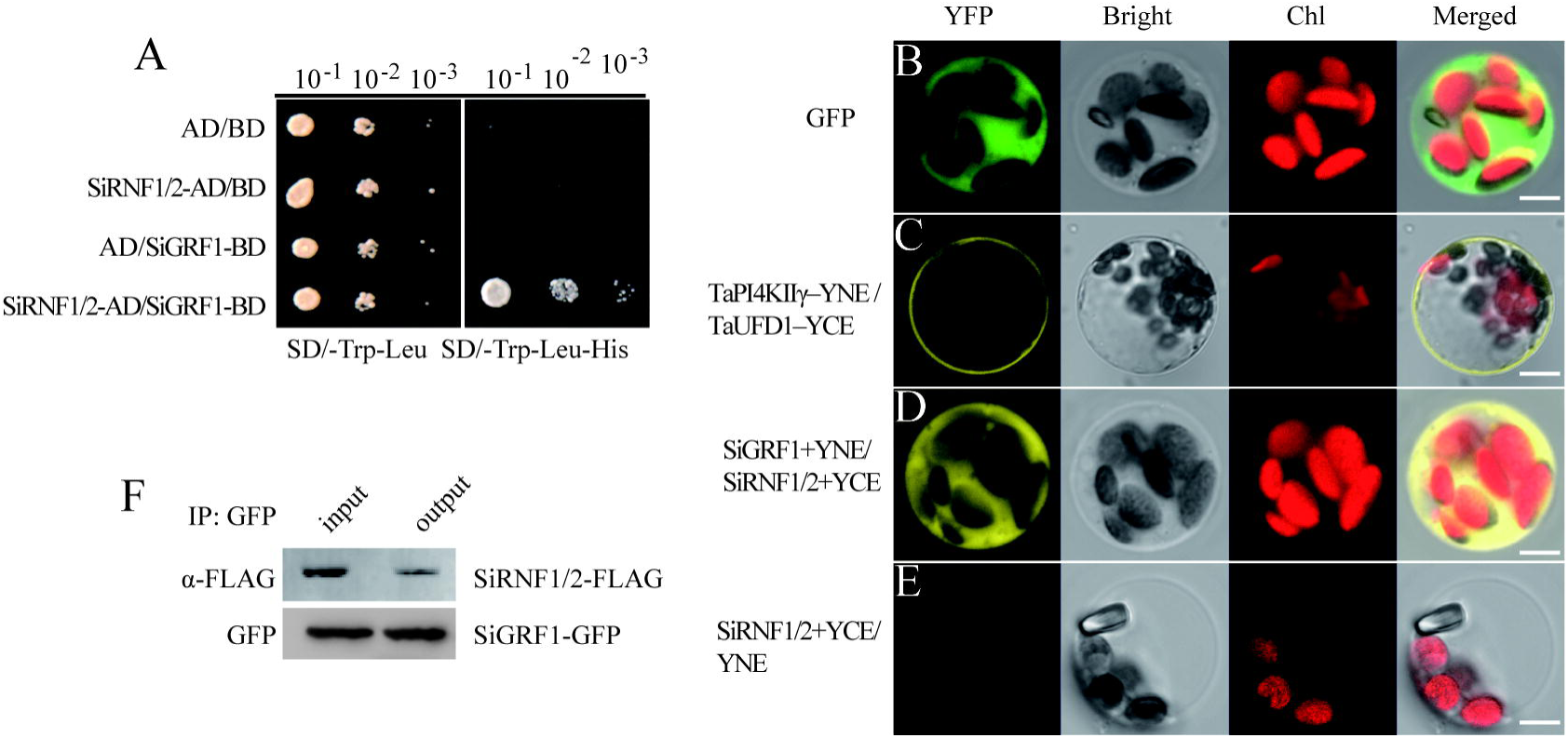
SiGRF1 interacted with SiRNF1/2 *in vivo*. (A) SiGRF1 interacted with SiRNF1/2 in yeast. Yeast strains expressing construct combinations of AD/BD, SiRNF1/2-AD/BD, AD/BD-SiGRF1 and SiRNF1/2-AD/BD-SiGRF1 were grown on media minus tryptophan and leucine (left panel) or minus tryptophan, leucine and histidine (right panel). (B) Green fluorescent protein localized throughout foxtail millet leaf protoplasts. (C) TaPI4KIIγ-YNE/TaUFD1-YCE was a positive control, TaPI4KIIγ and TaUFD1 can interact with each other in the plasma membrane. (D) Combinations of SiGRF1-YNE and SiRNF1/2-YCE were transiently co-expressed in foxtail millet leaf protoplasts. The YFP fluorescence indicates BiFC signals. Individual and merged images of YFP and Chl autofluorescence (Chl), as well as bright-field images of protoplasts are shown. Scale bars = 5 μm. (E) SiRNF1/2-YCE and YNE transiently co-expressed in foxtail millet leaf protoplasts. (F) SiGRF1-SiRNF1/2 interaction analysis using Co-IP. Anti-GFP affinity gels were used for immunoprecipitation and an anti-FLAG antibody was used to detect SiRNF1/2.

### Expression pattern and subcellular localization of SiGRF1

The *SiGRF1* gene has four introns and five exons with an open reading frame of 789 bp and putatively encodes a 31.4-kD 14-3-3 protein. To evaluate the expression pattern of *SiGRF1*, we performed qPCR using mRNA from different organs of foxtail millet, including root, culm, leaf, panicle, sheath and node (Figure 9 A). The results showed that *SiGRF1* was expressed in different tissues, and the expression of *SiGRF1* in panicles, leaves and node were higher than that in roots, culm, and leaf sheaths (Figure 9 A). To further characterize the expression pattern of *SiGRF1*, the GUS (β-glucuronidase) reporter gene driven by the 2.5-kb promoter region of the *SiGRF1* gene was introduced into *A. thaliana*. These transgenic lines were used for histochemical assays at different developmental stages (Figure 9 B-K). We observed strong GUS activity in the root apical meristem (RAM) and root cap columella (Figure 9 B), as well as the stele, lateral root primordium, and hypocotyl (Figure 9 C, D, and E). With the growth of plants, the *SiGRF1* gene was expressed in the ripening siliques, sepals, pollen grains, petiole, mesophyll and stoma (Figure 9 F-K). To explore the subcellular localization of the SiGRF1 protein, we transiently expressed the SiGRF1-GFP fusion gene in foxtail millet leaf protoplast. After an overnight incubation, the protoplasts were analyzed using a confocal microscope. We found that SiGRF1-GFP was localized only in the cytoplasm in foxtail millet protoplasts (Figure 9 L-O).

**Figure 9.**
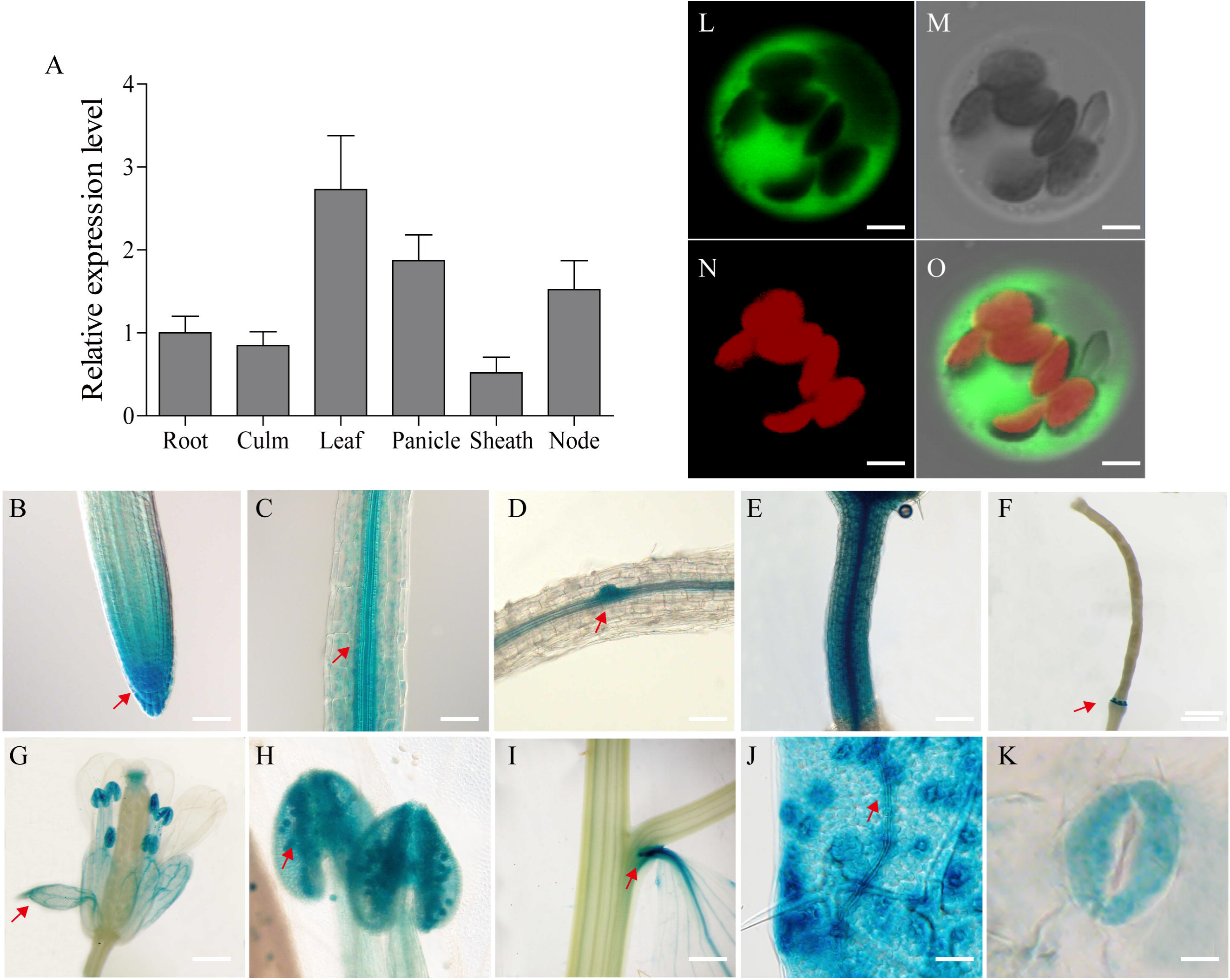
Tissue-specific expression of the *SiGRF1* gene and subcellular localization of its coding protein. (A) Analysis by RT-PCR of *SiGRF1* expression patterns. *SiActin* was used as the control. (B–K) Root apical meristem (RAM), root cap columella, stele, lateral root primordium, hypocotyl, ripening siliques, sepals, pollen grains, petiole, mesophyll and stoma. Scale bars = 500 μm. (L–O) SiGRF1 localized to the cytoplasm. Green fluorescence indicates GFP and red fluorescence indicates stained protoplasts. Bars = 5 μm.

## Discussion

Highly conserved in all eukaryotes, 14-3-3 proteins are phosphopeptide-binding proteins (Aitken et al., 1992). In this study, we identified eight 14-3-3 proteins in the foxtail millet genome. Our results agree with the research of Kumar, et al. (2015). The phylogenetic analysis of the 14-3-3s from seven species, including the eudicot *A. thaliana* and the monocots *B. distachyon*, *O. sativa*, *T. aestivum*, maize, *S. bicolor*, and foxtail millet, revealed that all of the 14-3-3 proteins can be categorized into ten discrete groups (Clusters I to □). Furthermore, there appeared to be a species-specific aggregation of genes in the seven species, as exhibited by groups □, □, and □ (Figure 1), which suggests that gene function may have species specificity, and these genes may have expanded after the separation of monocots and dicots.

The sequences of SiGRF2, SiGRF3 and SiGRF5 have high similarity and were categorized into Cluster □. The three genes were also found to be weakly expressed in total aerial tissues under blue light and far red light treatments (Figure 3). Together, the data suggest that these genes may have functional redundancy in regulating plant light response. In addition, the homologous genes of *SiGRF4* and *SiGRF7* were *T. aestivum TaWIN2* and *TaWIN1*, respectively. TaWIN1 and TaWIN2 can directly interact with WPK4, which is responsible for controlling the nitrogen metabolic pathway (Ikeda et al., 2000). We speculate that *SiGRF4* and *SiGRF7* may have similar functions to *TaWIN2* and *TaWIN1*.

Microarray analysis of the expression of the *SiGRF* genes in the four major foxtail millet tissues revealed that six of these genes were differentially expressed in at least one tissue. The tissue-specific expression profiling of *SiGRF*s could facilitate the combinatorial usage of *SiGRF*s in transcriptional regulation of different tissues, whereas ubiquitously expressed *SiGRF*s might regulate the transcription of a broad set of genes. Notably, *SiGRF1* and *SiGRF2* were strongly expressed only in the germ shoot but further investigation is needed to elucidate their potential functions. The low expression of *SiGRF5*, *SiGRF7* and *SiGRF8* observed in the leaf suggests that these three genes might be involved in the regulation of plant leaf development. Strong expression of *SiGRF6* in all plant tissues suggest that this gene may be involved in many physiological processes (Figure 2). In addition, overexpression of the *SiGRF* gene likely reduced sensitivity of transgenic plants to abiotic stress and exogenous ABA during seed germination and seedling growth. However, the stress response mechanisms of SiGRF need further study.

Current data clearly show plant 14-3-3 proteins are involved in many key physiological processes, ranging from metabolism to transport, biotic and abiotic stress responses, hormone signaling pathways (brassinosteroids, auxin, ABA, gibberellins, ethylene, cytokinins), plant growth and development, and flowering (Chen et al., 2006; Yuan et al., 2007; Barjaktarović et al., 2009; Yekti Asih et al., 2009; Xiaohua et al., 2010; Denison et al., 2011; Sun et al., 2015; Camoni et al., 2018). For example, two *A. thaliana* proteins, 14-3-3μ and 14-3-3υ, influence transition to flowering and early phytochrome response; loss of function of *14-3-3*μ and *14-3-3*υ showed a delay in flowering of 3–5 days under long-day conditions (Mayfield et al., 2007). In contrast, rice *GF14c* causes earlier flowering by 12–17 days, whereas overexpression of *GF14C* causes a 5–20 day delay in flowering under short day conditions (Yekti Asih et al., 2009). Additional protein interactions with 14-3-3s have been shown with CONSTANS, FT, PHOTOTROPIN1 and PHOTOTROPIN2 in *Arabidopsis*; FT orthologs of Heading Date 3A in rice ((Mayfield et al., 2007); and SELF-PRUNING in tomato (Pnueli et al., 2001). These protein interactions function in floral photoperiodism, blue light signaling and the switching between determinate and indeterminate plants.

Plants cannot move, so them has to face various environmental stresses during their life span, and flowering is crucial to successful reproduction in flowering plants. Therefore, in order to survive, plants have evolved a series of mechanisms to avoid, tolerate, or even resist hostile-environment stress signals to ensure reproduction (Levy and Dean, 1998). Previous studies have shown that pathogen infection, drought, and abnormal temperatures are able to accelerate flowering time (Korves and M., 2003; Meyre et al., 2004; Kumar et al., 2012; Xu et al., 2014). Salt stress is also considered as a negative factor on flowering time in most plants (Apse et al., 1999; Yu et al., 2017). Current research shows that salinity-delayed flowering is caused by a DELLA-dependent pathway. Salinity elevates the stability of DELLA proteins, which can restrain plant growth (Achard et al., 2006). In the quadruple-DELLA mutant, the response of salt-delayed flowering disappeared in plants under salt stress, and the expression level of *LFY* did not change under salt stress (Achard et al., 2006). Research shows NTM1-LIKE 8 mediates salinity-delayed flowering initiation via suppression of *FT* expression (Kim et al., 2007). In addition, *BROTHER OF FT AND TFL1* (*BFT*) overexpression in transgenic *A. thaliana* resulted in a delayed flowering phenotype when compared with that of the WT under normal or saline conditions. However, deletion of the *BFT* gene resulted in a normal flowering time in *bft-1* mutants exposed to high salinity (Ryu et al., 2011). The constitutive high level of expression of *WRKY71* can induce earlier flowering by promoting *FT* and *LFY* expression to act against the inhibition by DELLAs, BFT, etc. Overexpressing *WRKY71* plants showed an early-flowering phenotype under high salinity conditions (Yu et al., 2017).

In this paper, we found a foxtail millet 14-3-3 protein (SiGRF1) involved in flowering in plants under salt stress. The flowering initiation time of *SiGRF1*-OEs occurred earlier than that of Col-0 under high salinity conditions. Similarly, the expression of *WRKY71*, *FT*, *LFY* and *FUL* in *SiGRF1*-OEs was considerably higher than those in Col-0 under salinity-stressed conditions. The results suggest that the *SiGRF1* gene may regulate the initiation date of flowering in plants exposed to salt stress by up-regulating the transcription level of *WRKY71* to promote *FT* and *LFY* expression to act against the inhibition by DELLAs, BFT, etc. (Figure 10). Overall our findings suggest that SiGRF1 activity hastens flowering time, thereby providing a means for the plant to complete its life cycle and avoid further exposure to salt stress. Thus, we reveal a potential mechanism of plants to avoid environmental stresses.

**Figure 10.**
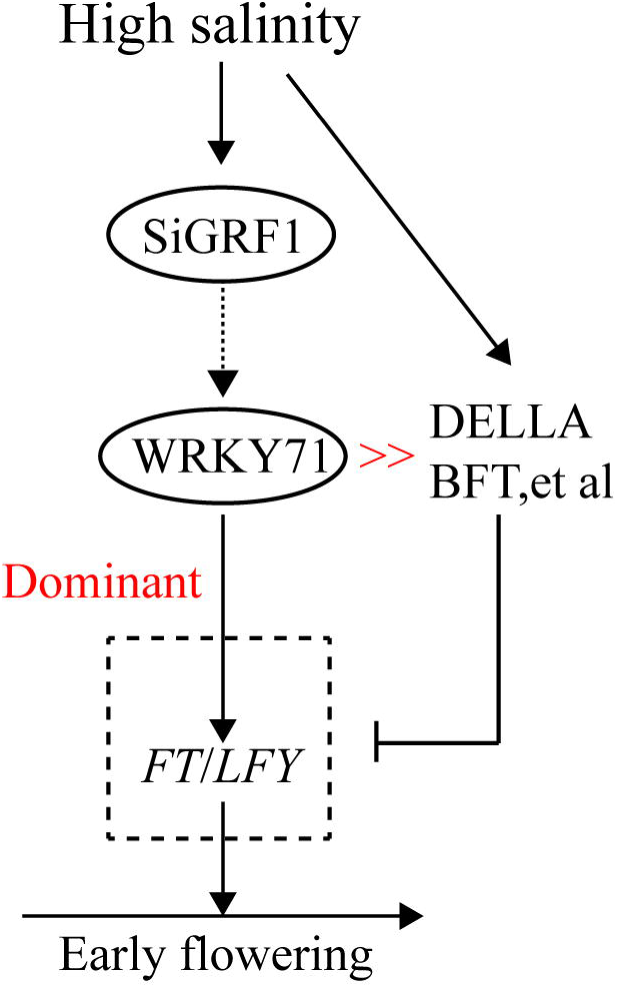
Proposed model of SiGRF1 function in flowering plants exposed to salinity treatment. High salinity promotes DELLAs, BFT, etc., and thereby inhibits *FT* and *LFY* expression. *SiGRF1* is induced by high salinity, which induces *WRKY71* to promote FT and LFY expression to act against their inhibition by DELLAs, BFT, etc. In the *SiGRF1*-OE plants under high salinity conditions, the highly elevated levels of gene expression of FT and LFY likely resulted in high levels of transcription. Thus SiGRF1-OE showed an early-flowering phenotype under high salinity conditions.

## Supplementary data

The following materials are available in the online version of this article.

**Table S1.** The sequences of primers used in the study. The sequences shown in lower case were added to generate a restriction enzyme site.

**Table S2**. *SiGRFs* in foxtail millet. Detailed genomic information including domain/class, alias, ORF length, protein length, genomic locus (chromosomal location), number of introns within ORF, subcellular localization, isoelectric point, and molecular weight (kDa) of the SiGRFs proteins for each *SiGRFs* gene.

**Table S3.** Average log signal values of SiGRF protein-encoding genes from three biological replicates of each sample.

**Table S4**. Average log signal values of SiGRF genes subjected to ammonia, drought, nitrate, urea, dark, blue, red, and far red light treatments.

**Figure S1.** The seed germination rates of *SiGRF1-*OEs, *SiGRF2-*OEs, *SiGRF3-*OEs, and *SiGRF4-*OEs under the no-stress and stress treatments.

**Figure S2.** The seed germination rates of *SiGRF5-*OEs, *SiGRF6-*OEs, *SiGRF7-*OEs, and *SiGRF8-*OEs under the no-stress and stress treatments.

**Figure S3.** Phenotypic comparison of root lengths of *SiGRF1-*OE, *SiGRF2-*OE, *SiGRF3-*OE, and *SiGRF4-*OE plants grown on MS medium with or without treatment of 120 mM NaCl, 6% PEG, or 0.5 μM ABA. Each treatment contained three independent replicates. Images were recorded on Day 5 after the transfer of 5-day-old seedlings from ½ MS medium to plates containing NaCl, PEG, or ABA. White solid line indicates that plants promote root growth. Bars = 1 cm. Data represent means ± SD (n = 30).

**Figure S4.** Phenotypic comparison of root lengths of *SiGRF5-*OEs, *SiGRF6-*OEs, *SiGRF7-*OEs, and *SiGRF8-*OEs plants grown on MS medium with or without treatment of 120 mM NaCl, 6% PEG, or 0.5 μM ABA. Each treatment contained three independent replicates. Images were recorded on Day 5 after the transfer of 5-day-old seedlings from ½ MS medium to plates containing NaCl, PEG, or ABA. White solid line indicates that plants promote root growth. Bars = 1 cm. Data represent means ± SD (n = 30).

## Authors’ contributions

WJZ conceived and coordinated the project; JML designed experiments, edited the manuscript, analyzed data, and wrote the first draft of the manuscript; CJJ analyzed data and performed experiments; LK and CCZ provided analytical tools and managed reagents; YS contributed valuable discussions. All authors have read and approved the final version of the manuscript.

## Acknowledgments

This research was financially supported by the National Natural Science Foundation of China (31871611 and 31960622) and China Postdoctoral Science Foundation (2018M643751).

## Competing interests

The authors declare they have no competing interests.

